# Robust division orientation of cambium stem cells requires cortical division zone components but not the preprophase band

**DOI:** 10.64898/2025.12.18.695170

**Authors:** Xiaomin Liu, Pantelis Livanos, Laura Sophie Schütz, Sabine Müller, Thomas Greb

## Abstract

The control of cell division plane orientation is fundamental for organizing developmental processes and shaping bodies of multicellular organisms. In plants, radial organ growth is mediated by the cambium, a stem cell niche deeply embedded in expanding organs and continuously producing xylem and phloem in a strictly bidirectional manner. Cambium stem cells (CSCs) are unique in comparison to other plant stem cells as they consistently divide along their longest axis clearly overriding the commonly found short axis rule. Despite the topological consistency of CSC divisions and the functional importance of proper division plane orientation for CSC derivatives, the regulatory mechanisms behind this consistency remain unknown. Here, we characterized microtubule organization during CSC divisions in *Arabidopsis thaliana* and found that division plane orientation is established independently from spindle orientation and the preprophase band (PPB). This conclusion is based on the remarkable variability of spindle orientation during metaphase and the robustness of CSC divisions in PPB-deficient mutants. Instead, the orientation of division planes depends on the cortical division zone (CDZ) and CDZ-related PHRAGMOPLAST ORIENTING KINESIN (POK) proteins. Our results highlight the importance of wood-forming CSCs and their subcellular characterization as an instructive example for the determination of cell division orientation in plants.

## Introduction

Cell division, a fundamental process administering growth and development in multicellular organisms, is precisely regulated in immobile plant cells. Often, plant cells divide along their minimal surface area and generate two equally sized daughter cells by following a ‘shortest axis’ division rule (Errera, 1888). However, divisions decisive for central developmental events frequently deviate from this rule. One example is radial organ growth in dicotyledonous species driven by CSCs, usually bi-directionally generating vascular tissues, xylem inwards and phloem outwards (Fischer *et al*., 2019; Shi *et al*., 2019; Smetana *et al*., 2019). CSCs are exceptionally elongated and divide along their longest axis clearly overriding the ‘shortest axis’ rule (Figure 1a). This property is assumed to be an essential functional trait because CSC derivatives differentiate into interconnected vascular elements whose elongated shape facilitates long distance transport of water, nutrients and organic molecules.

**Figure 1.**
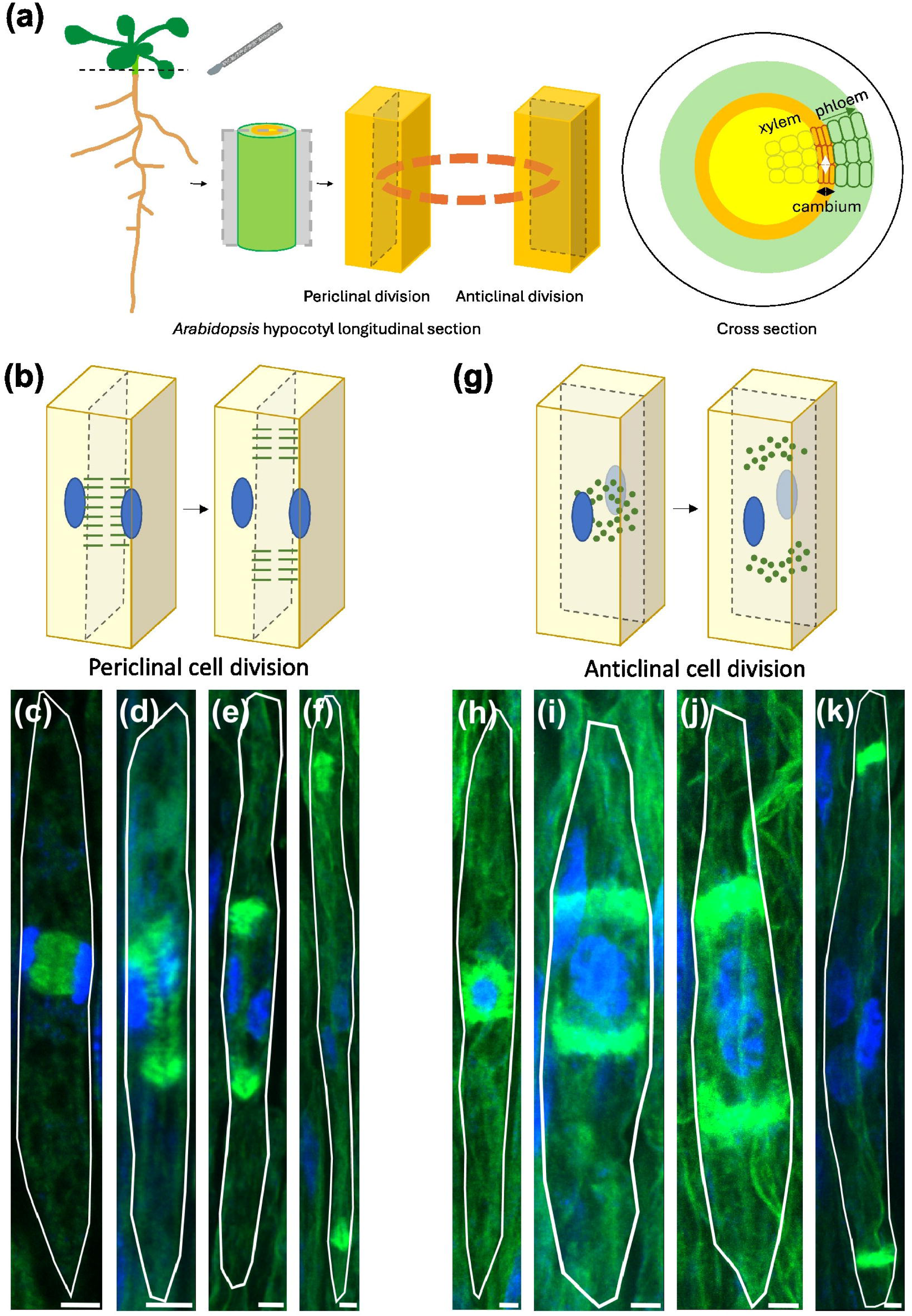
Determination of periclinal and anticlinal cell divisions by visualizing successive stages of phragmoplast microtubule organization after anaphase in dividing CSCs. (a) Scheme indicating periclinal and anticlinal cell divisions in CSCs in the Arabidopsis hypocotyl. The orange dashed ring in the middle panel represents the cambium area. The orange cuboids represent individual CSCs. The dashed panels represent longitudinal sections of hypocotyls or the future cell division planes in CSCs. The scheme on the right indicates CSCs (orange), periclinally generating xylem (yellow) or phloem cells (blue) and increasing the stem cell domain by anticlinal cell divisions. Black arrows indicate the direction of periclinal cell divisions, white arrows, of anticlinal cell divisions. (b-f) Phragmoplast microtubule organization in CSCs undergoing periclinal divisions. (g-k) Phragmoplast microtubule organization in CSCs undergoing anticlinal divisions. Phragmoplast microtubule arrays were detected by immunolabeling in CSCs using an antibody against tubulin (green). Nuclei were stained by DAPI (blue). (c, h) disk phragmoplast; (d, i) ring phragmoplast; (e, f, j, k) discontinuous phragmoplast. Cell boundaries are indicated by the white outline. n = 10 cells. Scale bar = 3 µm.

While consistently orienting along the longest cell axis, division planes in CSCs still have typically two possible orientations. Either they orient periclinally, i.e. parallel to the organ surface, thereby supporting the increase of the organ radius through the formation of xylem and phloem. Alternatively, they orient anticlinally, i.e. perpendicular to the organ surface, thereby increasing the circumferential dimension of the CSC domain (Figure 1a) (Tomescu & Groover, 2019). Regulatory factors influencing the remarkable orientation of these divisions and therefore plant vascular tissue organization are basically unknown. This is partly due to the fact that CSCs are deeply embedded in optically dense tissues preventing live imaging and direct physical manipulation. Recently, modelling of mechanical stress patterns together with mechanical perturbation in the radially growing *Arabidopsis thaliana* (Arabidopsis) hypocotyl indicated that CSCs experience mechanical stress generated by CSC-dependent tissue production and used as a stimulus to guide a periclinal orientation of division planes (Hofler *et al*., 2024). Moreover, it was hypothesized that the major stress direction within CSCs aligns with the longitudinal direction of the hypocotyl. This implies that the plane of maximal stress is perpendicular to the plane of minimal area and generally instructs cells to divide along their longest axis. These considerations align well with effects of more traditional mechanical perturbations on isolated cambium domains during which organized divisions could only be achieved when these domains were under external mechanical pressure (Lintilhac & Vesecky, 1984).

Four specialized microtubule arrays are characterized in plants: the cortical microtubule array, the PPB, the spindle, and the phragmoplast (Hamada, 2014). During interphase, microtubules are enriched at the cellular cortex, guiding structures and molecules like extracellular cellulose microfibrils laid down during cell wall synthesis to bear the load of turgor pressure and responding to external mechanical stress (Li *et al*., 2015). At the onset of cell division, during late G2, microtubules organize in PPBs marking the future division plane in the form of a cortical ring. The organization of cortical microtubule arrays, including the PPB during G2/prophase is accomplished by a protein phosphatase complex (PP2A) targeted to microtubules by TONNEAU1 RECRUITING MOTIF (TRM) proteins (Spinner *et al*., 2013). After the end of prophase, the bipolar spindle apparatus forms perpendicular to the PPB ring during prometaphase (Ambrose & Cyr, 2008) administering the accurate separation of the duplicated nuclear chromosomes in anaphase. Next, the phragmoplast forms between establishing daughter nuclei directing vesicles to the plane of cell division. There, vesicles fuse and expand the cell plate towards the cortical site previously defined by the PPB (Pickett-Heaps & Northcote, 1966).

The PPB is part of the so-called cortical division zone (CDZ), a polarized region corresponding to the selected cell division plane (Smertenko *et al*., 2017; Livanos & Muller, 2019). The PPB is known to facilitate the arrival of CDZ components (Smertenko *et al*., 2017; Livanos & Muller, 2019; Livanos *et al*., 2025). However, there is evidence that another function of the PPB is to limit spindle rotations in order to increase the robustness of cell division orientations. This conclusion is based on the observation that in *trm6;trm7;trm8* (*trm678*) triple mutants compromised in PPB formation, division planes are more variable. Nevertheless, PHRAGMOPLAST ORIENTING KINESIN1 (POK1) and POK2, two members of the kinesin-12 family in Arabidopsis, which co-localize with the PPB during prophase and function redundantly in maintaining the CDZ (Muller *et al*., 2006; Lipka *et al*., 2014) are still present in *trm678* cytokinetic cells (Schaefer *et al*., 2017; Livanos *et al*., 2025). Underlining POKs function, *pok1;pok2* double mutants show strongly disoriented cell division planes and a severely dwarfed phenotype (Muller *et al*., 2006), whereas *trm678* mutant phenotypes are considerably milder.

In this study, we characterized, for the first time, CSC divisions on the subcellular level and investigated how CSC’s division plane orientation is realized during mitosis. Through systematically determining phragmoplast orientation, we found that around 60 % of CSC divisions promote vascular tissue generation by periclinal cell divisions. During metaphase and anaphase, spindle arrays were considerably more variable than in other dividing cells (Schaefer *et al*., 2017), raising uncertainty about the role of PPBs during CSC divisions (Ambrose & Cyr, 2008). This uncertainty was further supported by observations that CSCs are devoid of PPB-related molecular makers and do not show altered division patterns in PPB-deficient mutants. In contrast, CSC division plane orientation strongly depended on CDZ components. Our results propose a model in which the robust orientation of CSC divisions along their longest axis depends on directional signals directly acting on CDZ orientation.

## Materials and Methods

### Plant material and plant growth conditions

In this study, the Col-0 accession of Arabidopsis was used. *trm678* mutants, *pok1;pok2* mutants and reporter lines of *TRM7-3xYFP*, *VHA-a3:VHA-a3-GFP*, *POK1:YFP-POK1* and *WOX4:mVenus-MBD* were described previously (Lipka *et al*., 2014; Schaefer *et al*., 2017; Smertenko *et al*., 2017; Lupanga *et al*., 2020; Hofler *et al*., 2024). Arabidopsis seeds were stratified at 4°C for two days in the dark before sowing. Seeds were sterilized with a few drops of 70 % EtOH and 0.05 % Triton X-100 by pipetting seeds up and down for one to two minutes, depending on the number of seeds. Then, the liquid was carefully poured out. 200 to 300 µl 100 % EtOH was added for one minute, and the liquid was poured out on filter paper under the clean bench. Sterilized seeds were sown on half MS solid medium. Plates were kept under short day conditions for one week. One-week-old seedlings were transferred from plates to soil and grown in growth chambers under short day conditions (10 h light, 14 h darkness, 22°C, 65 % humidity). Three-week-old seedlings were then transferred from short day to long day conditions (16 h light, 8 h darkness, 22°C, 50 % humidity).

### Immunolabelling

Immunolabelling of microtubules (α-Tubulin) was described previously (Zhao *et al*., 2020). During this study, we applied this protocol to visualize microtubule organization in CSCs with some modifications. The cell permeabilization buffer contained 1 % Driselase (w/v) and 1 % Triton X-100 (v/v) in microtubule stabilizing buffer (Du *et al*., 2021). Sections were incubated in the permeabilization buffer at 37°C for 20-25 min. For immunolabelling of tubulins, Monoclonal Anti-α-Tubulin antibody (Sigma-Aldrich, Cat# T5168) and Goat anti-Mouse IgG (H+L) Cross-Adsorbed Antibody, Alexa Fluor™ 488 (ThermoFischer Scientific, Cat# A-11001) were used as primary and secondary antibodies, respectively. For co-immunolabelling of tubulins and GFP-derived fluorescent proteins, besides antibodies mentioned above to visualize microtubules, GFP Polyclonal Antibody (ThermoFischer Scientific, Cat# A-11122) and Donkey anti-Rabbit IgG (H+L) Highly Cross-Adsorbed Secondary Antibody, Alexa Fluor Plus 594 (ThermoFischer Scientific, Cat# A-32745) were used. For immunolabelling of tubulin and other proteins using wild type, *trm678* mutants, *TRM7-3xYFP*, *pPOK1:YFP-POK1*, 37 to 39-day-old seedlings were used to generate longitudinal sections of hypocotyls.

### Histological analysis

Sample preparation and manipulation for histological analysis were performed as previously described (Wang *et al*., 2019; Wallner *et al*., 2020). Hypocotyls were collected in embedding cassettes and immersed in 70 % EtOH for at least two days. Samples were then processed using the LEICA ASP200S tissue processor and embedded in paraffin. The embedded samples were sectioned into 10 μm thick sections using the Leica microtome RM2235. After putting the section onto slides, they were stained by 0.05 % toluidine blue and mounted using Micromount Mounting Media (Cat# 3801731, Leica). The mounted slides were left at room temperature for more than two days. Finally, slides were scanned using the slide scanner Pannoramic SCAN II (3DHistech, Budapest, Hungary). Images were analyzed using the CaseViewer 2.2 software. For wild type and *trm678* mutants, cross sections were performed using 28-day-old seedlings, and longitudinal sections were performed using 38-day-old seedlings. For wild type and *pok1;pok2* mutants, cross sections were performed using 32-day-old seedlings which had not bolted yet. Longitudinal sections were performed using 42-day-old seedlings after bolting.

### Reporter analysis

Sample preparation and manipulation for reporter analysis were performed as previously described (Shi *et al*., 2019; Hofler *et al*., 2024). Briefly, hypocotyls of 38-day-old *VHA-a3:VHA-a3-GFP* were fixed overnight at 4°C by 4 % paraformaldehyde (PFA) and washed three times by phosphate buffered saline. For live-cell imaging of microtubules and YFP-POK1, 37-day-old seedlings of *WOX4:mVenus-MBD* and *pPOK1:YFP-POK1* lines were used. Hypocotyls were embedded in 7 % low melting agarose and free-hand longitudinal sections were generated using a razor blade. Sections were then placed in a two-well glass-bottom dish (ibidi, Cat# 80287) and immersed in water.

### Quantification of spindle and phragmoplast orientation

To analyze the orientation of spindles and/or the phragmoplast in dividing cambium cells of wild type and *trm678* mutants, immunolabelling of microtubules and staining of nuclei by DAPI (Sigma-Aldrich, Cat #D9542) were used. Image analysis was conducted using FIJI (Schindelin *et al*., 2012). In individual dividing CSCs, spindle angles were defined as the angle of aligned chromosomes relative to the longest CSC axis. 0° represented a spindle whose pole-to-pole axis was perpendicular to the longest axis of cambium cells. Phragmoplast angles were defined as the angle enclosed by aligned metaphase chromosomes and the longest CSC axis. Thus, 90° represented a spindle whose pole-to-pole axis was parallel to the longest axis of cambium cells. Consistently, 0° indicating a parallel alignment of the antiparallel microtubules relative to the CSCs’ longest axis. Quantification of spindle angles and phragmoplast angles was performed using the ‘angle tools’ in FIJI.

### Confocal microscopy

A Stellaris 8 confocal microscope and a 20x air objective (HC PL APO CS2, 20X/0.75 DRY) and a 63x water objective (HC PL APO CS2, 63x/1.2 WATER) were used in this study. The Diode 405 nm and WLL lasers were used for excitation. Image acquisition was performed using a 1024×1024 pixel scanning field, 4x line accumulation, pinhole 1AU and 400 Hz scanning speed. For (co-)immunolabelling experiments visualizing microtubules (and other proteins) together with nuclei or chromosomes in dividing CSCs, DAPI, tubulin and YFP/GFP fusion proteins were excited at 405 nm, 514 nm and 590 nm and detected at 425-494 nm, 519-576 nm and 600-672 nm, respectively.

### Single cell transcriptome analysis

For single cell transcriptome analysis, a previously published dataset (Zhao *et al*., 2025) eposited at Gene Expression Omnibus under accession code GSE224928 was analyzed using the Seurat library of R (Hao *et al*., 2021). Cells with nCount_RNA between 1201 and 9999 and a mitochondrial genome read fraction of less than 20 % were kept for further analysis. Clustering and UMAP were generated using default settings of Seurat with the parameters “dims = 1:15, resolution =1.2”. Marker genes for each cluster were generated by using the “FindAllMarkers” function of Seurat (only.pos = TRUE, min.pct = 0.25, logfc.threshold = 0.25). Individual genes were displayed via the “FeaturePlot” function. The code used for data analysis is found at the GitHub repository [https://github.com/thomasgreb/Liu-et-al_CSC-divisions].

## Results

### Phragmoplast organization reflects CSC division orientation

To investigate the orientation of division planes in CSCs during cytokinesis, we initially visualized successive stages of phragmoplast organization by immunolabeling microtubules in dividing cells. The phragmoplast is organized as a bipolar array of antiparallel microtubules that aid cell plate synthesis at the plane of cell division (Smertenko *et al*., 2017). Usually, the phragmoplast initially forms a disk of microtubules and expands towards the cortex by microtubule turn over. During expansion, the disk then transforms into a ring that becomes discontinuous upon fusion of the cell plate with the parental cell wall (Smertenko *et al*., 2017). Throughout the different stages, phragmoplast organization and nucleus topology typically reveal the future cell division orientation which we first used to distinguish periclinal from anticlinal CSC divisions (Figure 1a). As periclinal cell divisions, we categorized those cases in which two nuclei were consistently observed in the same optical plane at both sides of the phragmoplast in radial orientation (Movie S1). Disk or ring phragmoplasts appeared in these cases as continuous bipolar structure, separated by a dark line corresponding to the developing cell plate, localized between the two nuclei (Figure 1b-d). Discontinuous phragmoplasts had expanded away from the cell center and were found close to both the apical and basal cell boundaries (Figure 1e, f). In contrast to periclinally dividing cells, CSCs undergoing anticlinal cell divisions were characterized by two nuclei present in different optical planes (Movie S2). In these cases, the disk phragmoplast had a circular shape, whereas later, phragmoplasts were visible as complete or discontinuous rings (Figure 1g-k). In total, we found that approximately 60 % of CSC divisions were in periclinal, while around 40 % were in anticlinal orientation, among 407 dividing cells analyzed (Table 1). These results indicated that, as proposed before (Spicer & Groover, 2010), CSCs predominantly promote the formation of vascular tissues by conducting periclinal cell divisions, and to a lower extend divide anticlinally increasing the stem cell pool in circumferential orientation. The orientation of the disk phragmoplast in periclinally dividing CSCs appeared slightly variable (Supplemental Figure 1) which was not possible to be observed in anticlinally dividing cells because, here, the phragmoplast oriented parallel to the optical plane. Besides the disk stage, the orientation of the phragmoplast microtubule arrays in dividing CSCs perfectly indicated the final division orientation along the longest cell axis in periclinal or anticlinal orientation. In general, we concluded that the final division orientation of CSCs is already reflected by the orientation of the phragmoplast during early cytokinesis.

**Table 1.**
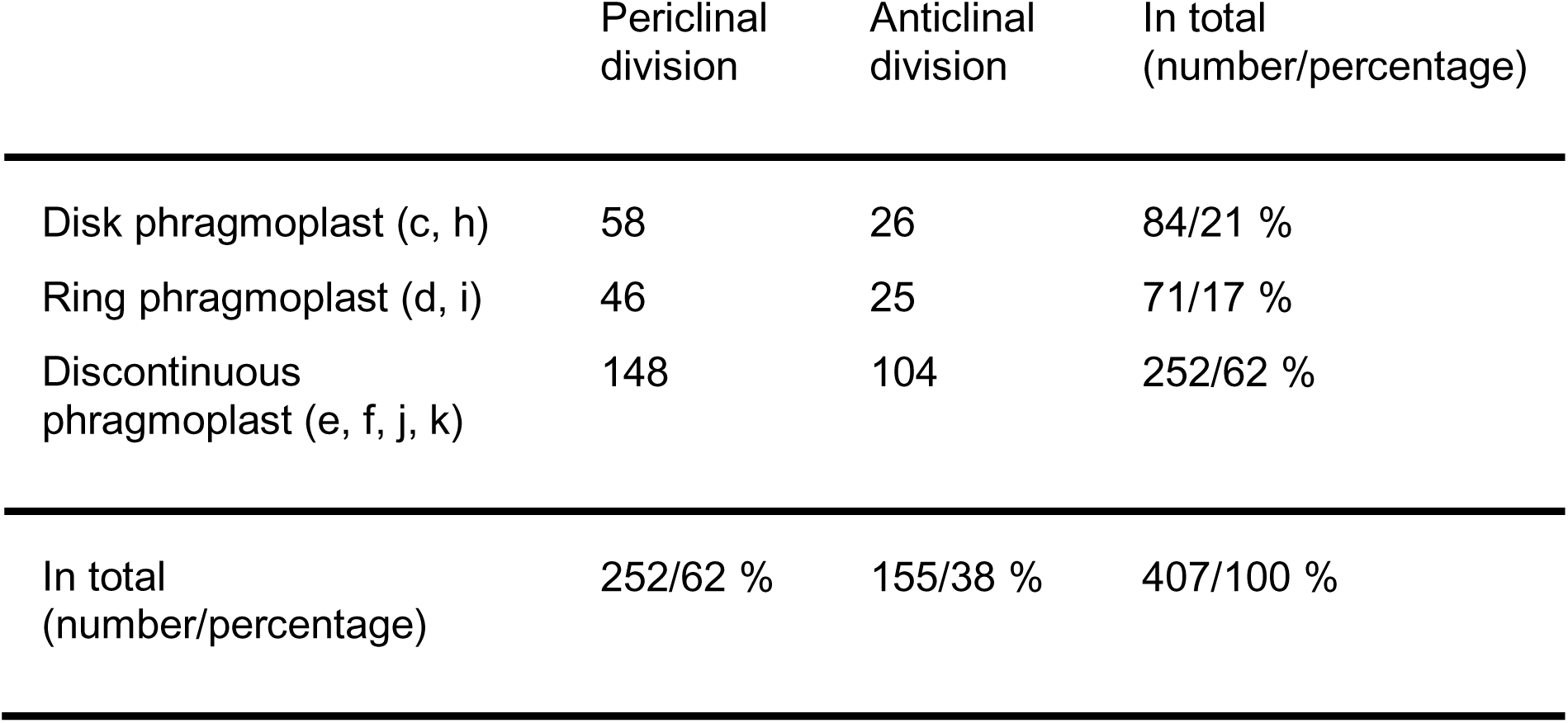
Quantification of the number and percentage of dividing CSCs visualizing three distinct phragmoplast microtubule arrays and distinguishing between periclinal or anticlinal divisions.

|  | Periclinal<br>division | Anticlinal<br>division | In total<br>(number/percentage) |
| --- | --- | --- | --- |
| Disk phragmoplast (c, h) | 58 | 26 | 84/21 % |
| Ring phragmoplast (d, i) | 46 | 25 | 71/17 % |
| Discontinuous<br>phragmoplast (e, f, j, k) | 148 | 104 | 252/62 % |
| In total<br>(number/percentage) | 252/62 % | 155/38 % | 407/100 % |

To calculate the relative duration of cell division stages, we further quantified the proportion of CSCs displaying the three distinct phragmoplast microtubule phases (disk, ring, and discontinuous). These analyses showed that the largest fraction of dividing CSCs (62 %) carried a discontinuous phragmoplast, representing the process of microtubule progression from the cell center to the apical and basal cross walls. The smallest fraction (17 %) of dividing CSCs showed a ring phragmoplast, which represented the beginning of microtubule array progression from the cell center to the poles and 21 % of dividing cells each showed a disk phragmoplast, respectively (Table 1). Thus, among different phragmoplast stages in CSCs, the progression of the phragmoplast toward the apical and basal cell boundaries appeared to take the longest. In comparison to other dividing plant stem cells, in which the phragmoplast only needs to span a rather short distance from the cell center to the opposite sides of the parental cell, phragmoplast expansion along the longest CSC axis, which typically extends to 50 - 100 µm, may therefore result in an overall prolonged cytokinesis. The latter conclusion was consistent with the notion that cytokinesis duration depends on the width of the cell (Gorst *et al*., 1986).

### Spindle orientation is highly flexible in dividing CSCs

Because the orientation of cell divisions along the longitudinal cell axis appeared already set during cytokinesis, we asked whether the same holds true during mitosis. In animal cells, spindle orientation is predominantly regulated by centrosomes and known to be dynamic (Bergstralh *et al*., 2017). In comparison, plant cells lack centrosomes and, determined by the PPB prior to mitosis, spindle orientation in dividing plant cells is usually very robust (Ambrose & Cyr, 2008). To reveal whether division orientation is already fixed in CSCs during mitosis, we determined spindle orientation during meta- and anaphase of cells undergoing periclinal divisions (Movie S3). CSCs undergoing anticlinal divisions were excluded from the analysis as the orientation of the spindle perpendicular to the optical plane prevented determining their angles (Movie S4). Angles of spindle/chromosome orientation in periclinally dividing cells were measured in relation to the longitudinal CSC axis, with 0° indicating that the orientation of chromosomes was parallel to their longest axis (Figure 2a-c). Interestingly, these measurements revealed a broad variation of orientations covering a range from 0° to 90°, suggesting that the final cell division orientation along the longest cell axis is independent from the orientation of the spindle. A weak maximum of orientations was observed around 45°, possibly due to a slight bias of this orientation caused by space constraints in the elongated CSCs (Figure 2f). In comparison to the disk phragmoplast whose orientation rotated within 20° in late division stages, the spindle orientation was therefore considerably more variable.

**Figure 2.**
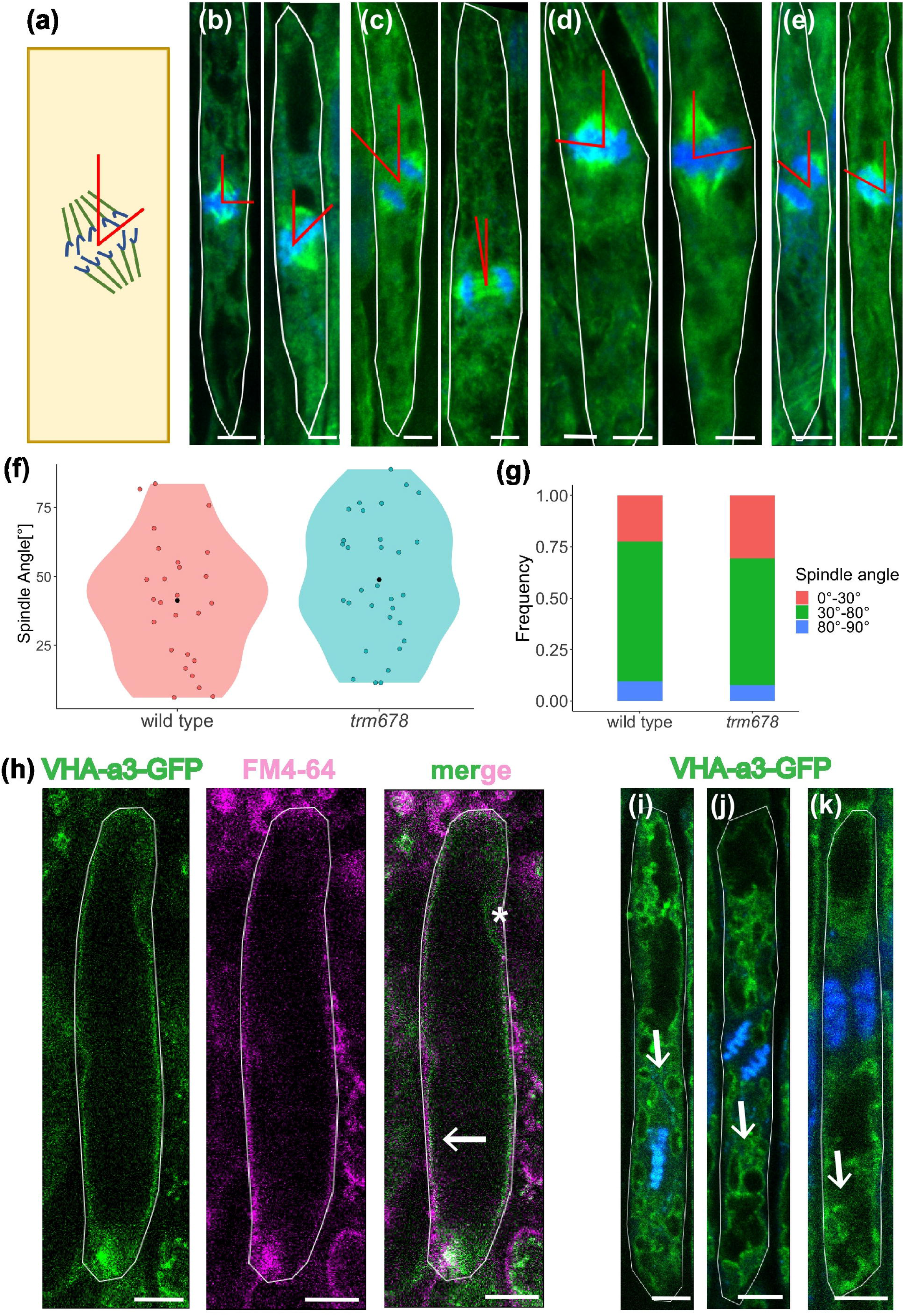
Initial spindle angles in CSCs are flexible in both wild type and *trm678* mutants. (a-e) Scheme of spindle orientation quantification in CSCs. Spindle angles (red) were measured by FIJI. Immunolabeling in CSCs was performed using an antibody against tubulin (green). Nuclei were stained by DAPI (blue). Cell boundaries are indicated by the white outline. (b,c) wild type. (d, e) *trm678* mutant. (b, d) metaphase spindle. (c, e) anaphase spindle. Scale bar = 3 µm. (f, g) Quantification of spindle angles in wild type (n = 26 cells) and *trm678* mutants (n = 31 cells). (f) violin plot. The mean is indicated as black dots. Individual data points are shown as pink or green dots. (g) boxplot. (h) Morphology of the vacuole in non-dividing CSCs was detected by live-cell imaging. The arrow indicates the overlap of vacuole (green) and the plasma membrane-derived signals (magenta) with the exception of the nucleus (asterisk). Cell boundaries are indicated by the white outline. n = 9 cells. Scale bar = 10 µm. (i-k) In dividing CSCs, the morphology of the vacuole was detected after PFA fixation. Nuclei or DNA were stained by DAPI (blue). Vacuole vesicles (green, VHA-a3) in metaphase (i), anaphase (j) or cytokinesis (k) are indicated by arrows. n = 3-5 cells. Scale bar = 5 µm. *VHA-a3:VHA-a3-GFP* was used in (h-k). Cell boundaries are indicated by the white outline.

Of note, the vacuole occupied most of the space in non-dividing CSCs, but changed significantly in organization, volume and size during divisions. Vacuolar distribution is essential for asymmetric cell division of the Arabidopsis zygote and is discussed to influence division plane orientation in other cell types (Panteris *et al*., 2004; Kimata *et al*., 2019). Visualizing vacuole morphology by the *pVHA-a3:VHA-a3-GREEN FLUORESCENT PROTEIN (GFP)* marker (Lupanga *et al*., 2020) and the plasma membrane by FM4-64 staining in live-cell imaging, revealed one central vacuole whose boundaries almost completely overlapped with the plasma membrane-derived FM4-64 signal in non-dividing cells (Figure 2h). For investigating vacuole morphology in dividing CSCs, we co-visualized vacuoles and chromosomes after PFA fixation again using the *pVHA-a3:VHA-a3-GFP* marker line. Instead of displaying an intact structure, the vacuole had fragmented into small vesicles during metaphase, anaphase or cytokinesis (Figure 2i-k). We speculate that this compartmentalization of the vacuole during mitosis is a necessary rearrangement that creates sufficient space for microtubule reorganization, chromosome alignment and segregation in dividing CSCs. In addition, it may allow for the observed variability of spindle orientation.

### Regular cell divisions in cambium stem cells do not rely on PPBs

In light of a rather narrow variation of spindle angles between 0°- 30° during mitosis in root tip cells (Schaefer *et al*., 2017) and of the strict orientation of the final CSC divisions along their longest axis, the broad range of spindle angle variation in dividing CSCs between 0°- 90° was surprising (Figure 2f). In this context it is important to note that the PPB is viewed as a stabilizing module aligning to predefined spatial cues, providing a sturdy cortical reference for the orientation of the spindle and the subsequent cell division (Schaefer *et al*., 2017). This feature of the PPB thus prompted us to investigate the role of the PPB in stabilizing the orientation of CSC divisions.

The TRM7 protein serves as a PPB indicator as it colocalizes with PPB microtubules forming a dual, cytoplasmic and cortical ring during late G2 or prophase (Schaefer *et al*., 2017). To visualize TRM7 during cell division we used a TRM7-3xYELLOW FLUORESCENT PROTEIN (TRM7-3xYFP) fusion protein expressed under the control of its endogenous promoter (Menges *et al*., 2003; Schaefer *et al*., 2017). Demonstrating the utility of the TRM7-3xYFP marker, we detected the PPB in dividing root tip cells by co-immunolabeling both microtubules and the YFP-tagged TRM7 protein. In these cells, TRM7-3xYFP was localized in both cytoplasmic and cortical areas, where it co-localized with PPB microtubules (Supplemental Figure 2a). However, when conducting co-immunolabeling of microtubules and TRM7-3xYFP in CSCs, we found that the TRM7-3xYFP protein was in five mitotic cells analyzed exclusively present in the cytoplasm and that there was neither a sign of a PPB-like microtubule formation nor a helical cortical array, found in CSCs during interphase (Supplemental Figure 2b). In contrast, we observed a high density of perinuclear microtubules identifying these cells as prophase CSCs (Figure 3, Movie S5). Interestingly, this conspicuous microtubule array in dividing CSCs closely resembled the situation of root tip cells in *trm678* mutants in which cells usually do not form a PPB but show a similar assembly of aberrant perinuclear microtubules (Schaefer *et al*., 2017). As already suggested previously, based on ultrastructure analyses (Evert & Deshpande, 1970; Farrar & Evert, 1997; Oribe *et al*., 2001), these observations indicated that CSCs do not form a PPB during their divisions.

**Figure 3.**
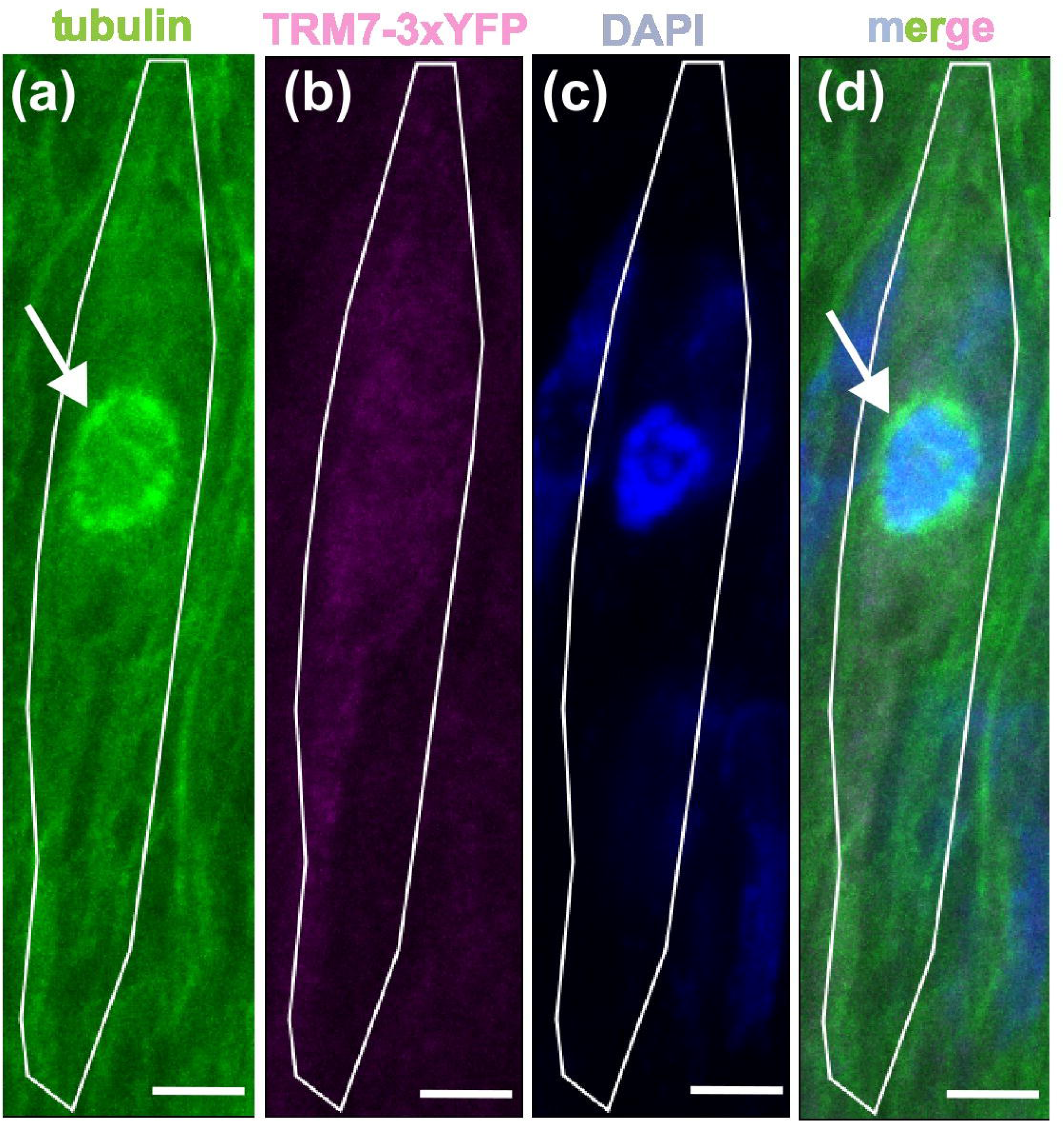
Perinuclear microtubule arrays are detected in dividing CSCs. (a-d) Maximum intensity projections of signals from perinuclear microtubule arrays detected by co-immunolabeling of tubulin and TRM7 in a dividing CSC using an antibody against tubulin (a, green) and an antibody against the YFP protein (b, magenta). Nuclei were stained by DAPI (c, blue). Cell boundaries are indicated by the white outline. Arrows point to the perinuclear microtubule arrays. The *TRM7-3xYFP* marker line was used in all analyses. n = 5 cells. Scale bar = 3 µm.

### Division orientation in the cork cambium is PPB-dependent

Radial growth is driven by two concentric bifacial meristems: the vascular cambium and, at the organ periphery, the phellogen. The activity of the phellogen, also known as the cork cambium, generates phellem (cork) outwards and phelloderm (cork parenchyma) inwards. The phellem-phellogen-phelloderm complex is collectively called the periderm, which forms the outer bark which is most prominent in woody species (Campilho *et al*., 2020). Given the role of the PPB in predicting cell division plane orientations, we performed histological analyses of hypocotyl cross and longitudinal sections in both wild type and PPB-deficient *trm678* mutants to reveal the functional significance of a PPB absence in CSCs and, as a functional comparison, in the phellogen.

When analyzing hypocotyl cross sections, we found the overall hypocotyl area to be decreased in *trm678* compared to wild type (Figure 4a-c), possibly reflecting an overall reduced growth of the *trm678* mutant (Schaefer *et al*., 2017). Importantly, *trm678* plants showed a normal cambium anatomy with highly regular cell division planes, whereas cell arrangement was much more irregular in the *trm678* periderm than in wild type (Figure 4d-g, Supplemental Figure 3). Especially, the generation of ordered files of compact cells derived from the phellogen was severely disturbed in *trm678* mutants, unlike the formation of regular cell files in the CSC cambium area (Figure 4e, g). This observation suggested that, during radial growth, PPB-deficiency interferes with cell division plane orientation in phellogen stem cells but not in CSCs. Because previous studies established a role of the PPB in limiting the rotation of the mitotic spindle, we compared spindle orientations between wild type and PPB-compromised *trm678* mutants. CSCs displayed the same variation in spindle rotation during metaphase and anaphase in both wide type and *trm678* mutants (Figure 2a-f). The frequency of CSCs showing spindle angles ranging from 0°-30°, 30°-80°, 80°-90° were similar between wild type and *trm678* mutants with most spindle angles ranging between 30° and 80° in both wild type and *trm678* mutants (Figure 2g). As mentioned before, such a broad variation of rotation angles was not observed previously in root tip cells of wild type plants, which showed variation within 30°, but was similar to the variation of spindle angles in root tips of *trm678* plants (Schaefer *et al*., 2017). Collectively, these results indicated that robust CSC division plane positioning is independent from PPB formation.

**Figure 4.**
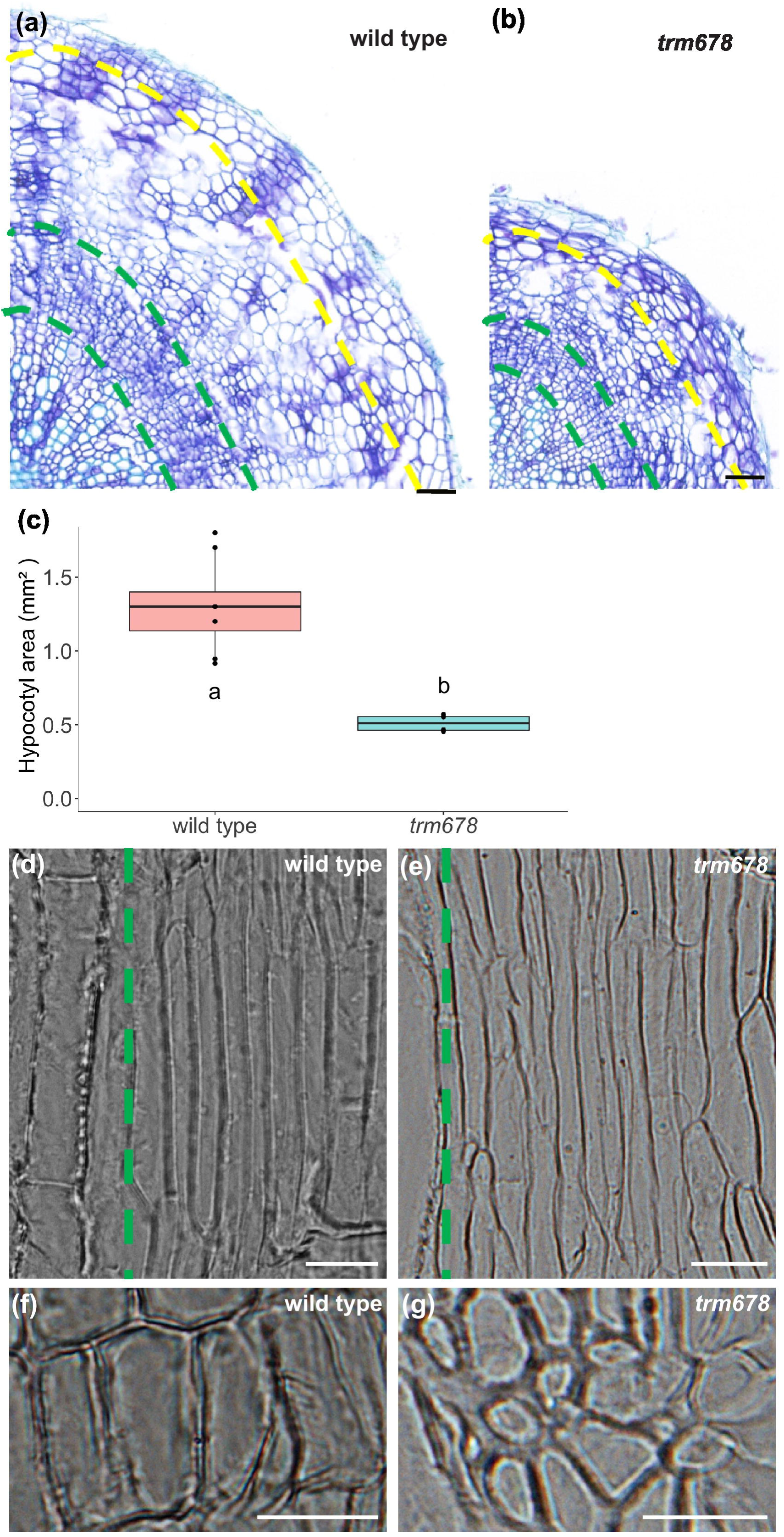
Histological analysis of wide type and *trm678* triple mutants. (a, b) Toluidine bule-stained hypocotyl cross sections from wild type (a) and trm678 mutants (b) embedded in paraffin. Scale bar = 50 µm. (c) Quantification of hypocotyl area in wild type and *trm678* mutants. Individual data points are shown as black dots. One-way ANOVA, n = 8 independent seedlings. P<0.0001. (d-g) Bright field analysis of longitudinal hypocotyl sections of wild type (d, f) and *trm678* mutants (e, g) embedded in Technovit 7100. (d, e) Cambium cell files are seen on the right side of the green dashed lines. The xylem is found on the left side. (f, g) Cell organization in the periderm. n = 5-8 independent seedlings. Scale bar = 20 µm.

### Orientation of CSC divisions depends on CDZ components

After having found indications that CSC divisions orient independently from the PPB, we investigated the role of the CDZ in determining CSCs’ division orientation. As mentioned, POK1 and POK2 proteins are universally required for the guidance of the phragmoplast towards a preformed CDZ to secure proper division plane orientation (Müller *et al*., 2006) and localize to the CDZ throughout cell division (Lipka *et al*., 2014; Herrmann *et al*., 2018). Indeed, when performing live-cell imaging in a *pPOK1:YFP-POK1* reporter line expressing POK1 as a fusion protein to YFP (Lipka *et al*., 2014), we found pairs of focused spots of YFP-POK1 at the apical and basal ends of CSCs (Figure 5a). To correlate POK1 localization with different stages of CSC divisions, we next performed co-immunolabeling of tubulin and the YFP-POK1 fusion protein using the same *pPOK1:YFP-POK1* line. In CSCs undergoing cytokinesis and showing, among others, a discontinuous phragmoplast microtubule array we observed the characteristic POK1 localization at opposing cell ends in images of median cell planes, as part of its ring-shaped assembly at the cortical division zone (Figure 5b, Lipka *et al*., 2014). These observations suggested that the CDZ is formed and establishes the position of division planes in dividing CSCs without forming a PPB. Consistently, *pok1;pok2* double mutants showed a dramatically disorganized cambium patterning where the typical concentric cambium organization had disappeared (Figure 6). In fact, in *pok1;pok2* plants it was not possible to distinguish CSCs from phloem cells, and cell morphologies were substantially altered (Figure 6). Therefore, *POK1* and *POK2* are necessary for cell division plane positioning and instruct tissue organization during radial plant growth.

**Figure 5.**
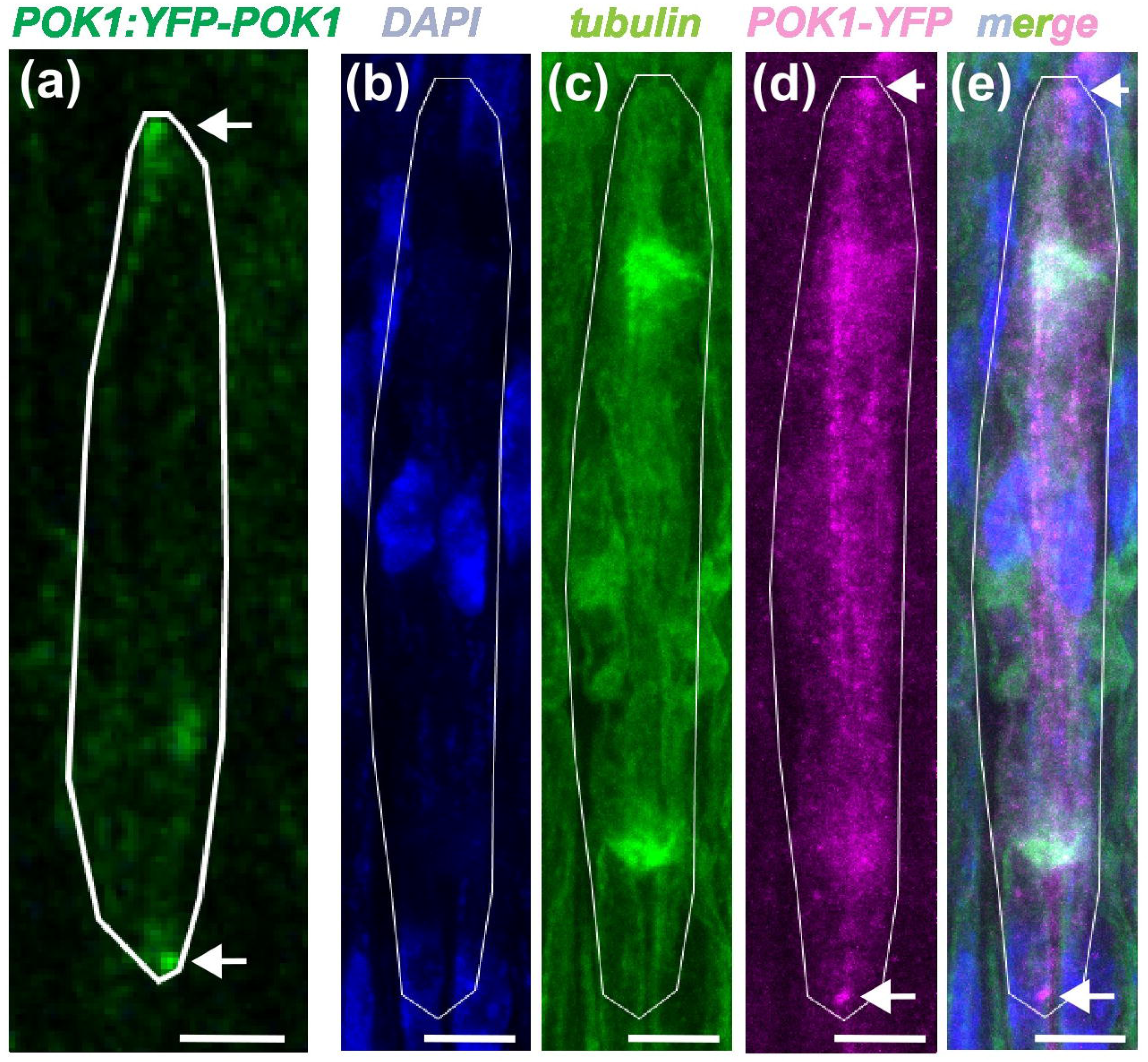
POK1 locates at opposite CSC poles during cytokinesis. (a) POK1 localized specifically at opposite CSC poles in longitudinal hypocotyl sections. n = 10 cells. Scale bar = 10 µm. (b-e) Maximum intensity projections of fluorescent signals during CSC cytokinesis detected by co-immunolabeling using an antibody against tubulin (green) and an antibody against the YFP protein (magenta). Nuclei were stained by DAPI (blue). n = 8 cells. Scale bar = 3 µm. The *pPOK1:YFP-POK1* line was used in all analyses. Arrows indicate POK1 signals at opposite cell poles. Cell boundaries are indicated by the white outline.

**Figure 6.**
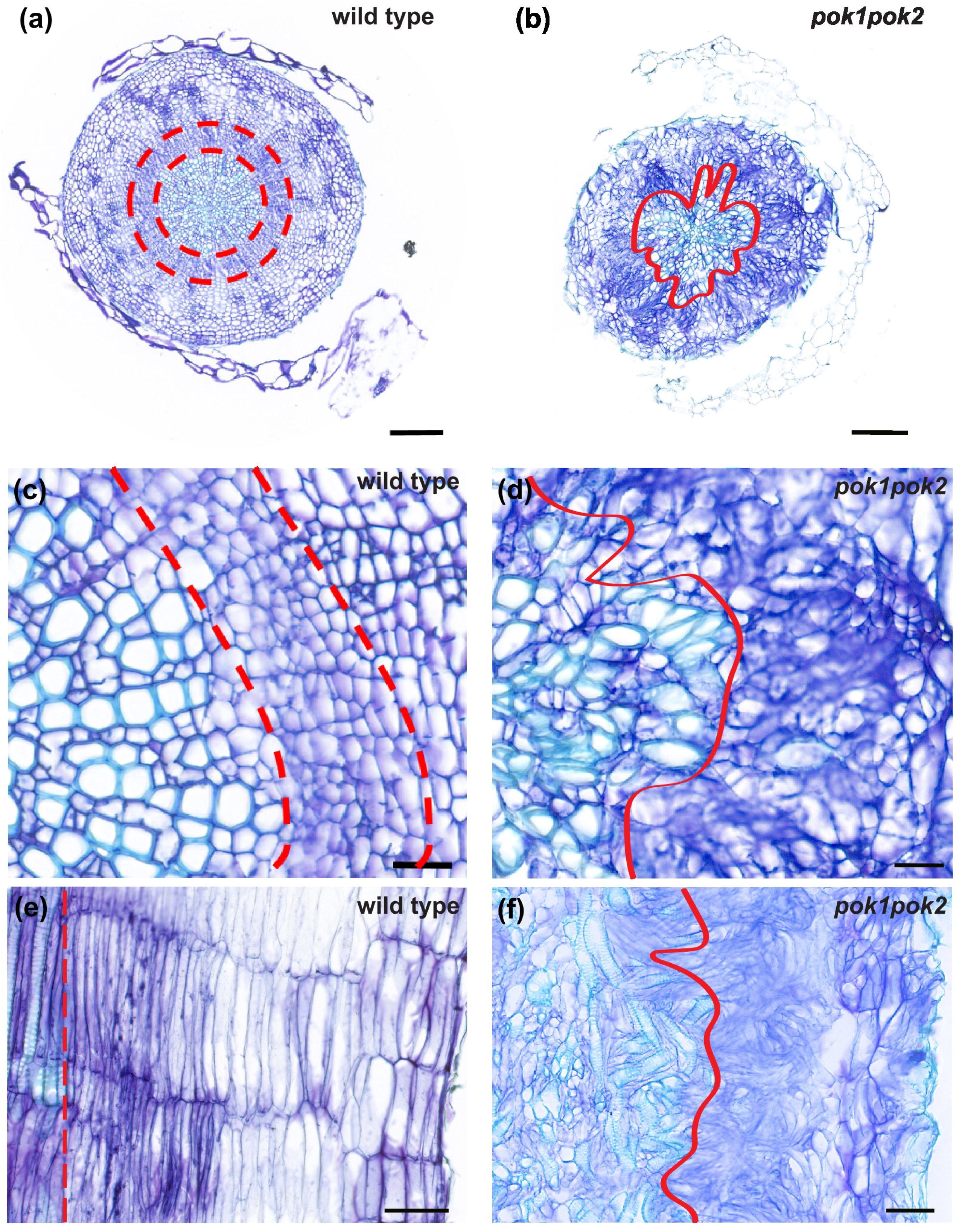
*pok1;pok2* double mutants show highly disorganized vascular tissues. (a, c, e) Hypocotyl cross (a, c) and longitudinal (e) sections of wild type stained by toluidine blue. (b, d, f) Hypocotyl cross (b, d) and longitudinal (f) sections of *pok1;pok2* double mutants stained by toluidine blue. The red dashed lines indicate the boundaries of the cambium. The continuous red line indicates the boundaries of the xylem in *pok1;pok2* double mutants. (c, d) are magnifications of (a, b). n = 5 independent seedlings. Scale bar (a, c) = 100 µm; Scale bar (b, d) = 20 µm. Scale bar (e, f) = 50 µm.

In line with a differential regulation of CDZ vs. PPB components, when analyzing single cell transcriptome data of the Arabidopsis hypocotyl published previously (Zhao *et al*., 2025), we found that the CDZ components POK1 and POK2 are expressed in CSCs and, especially, in dividing hypocotyl cells (Supplemental Figure 4). In contrast, genes encoding the protein phosphatase scaffolding subunit PP2A1 and TRM8, both involved in PPB formation, showed a rather unspecific activity pattern throughout hypocotyl tissues (Supplemental Figure 4). This demonstrated that, in the hypocotyl, regulation of PPB assembly is uncoupled from CDZ formation on the transcriptional level.

## Discussion

Oriented cell divisions generate cell diversity and shape tissues in multicellular organisms. Compared to plant cells, animal cells lack cell walls to maintain their cell shape during growth and development and experience dramatic changes in morphology during mitosis. Typically, dividing animal cells form spheres and remodel the actomyosin cortex for normal bipolar spindle formation (Stewart *et al*., 2011; Lancaster *et al*., 2013). Importantly, the position and orientation of the mitotic spindle determines the orientation of the cleavage furrow and, thus, contribute to the relative size and arrangement of daughter cells (Rappaport & Rappaport, 1974). In comparison, the PPB and the CDZ predict and preserve division plane orientation in plant cells, while spindle orientation usually does not have a strong effect on the placement of the new cell plate (Livanos & Muller, 2019).

The cambium is essential for radial plant growth and executes highly ordered divisions in CSCs. Unlike other plant stem cells, CSCs show a considerable degree of cellular differentiation and divide exclusively along their longest axis, deviating substantially from the ‘shortest axis’ division rule. Considering their unique morphology, we propose that it is instructive to characterize mitosis in CSCs to address the question of how cell division planes are determined in general, a question which has basically not been fully answered in plants. However, CSCs are embedded deeply in optically dense tissues, bringing them out of reach for being observed in intact organs and making them only accessible visually after transverse or longitudinal sectioning. This makes it exceptionally difficult to reveal the dynamics of their subcellular structures during mitosis.

Consistent with previous reports, that CSCs do not form PPBs (Evert & Deshpande, 1970; Farrar & Evert, 1997; Oribe *et al*., 2001), in this study, we found that the PPB reporter protein *TRM7-3xYFP* did not localize to polarized cortical regions of the cell and we did not find indications for a PPB-like microtubule assembly in dividing CSCs. Instead, CSCs showed an enriched perinuclear microtubule array as being found in dividing root cells of PPB-deficient *trm678* mutants. Moreover, *trm678* mutants did neither show alterations in cambium anatomy nor in CSC’s spindle orientation. Taken together, these observations indicate that, in spite of their highly ordered cell divisions, there is no relevant function of the PPB in CSCs. In addition to CSC divisions, endosperm cellularization and meiocyte development are examples of processes in which cells undergo well-ordered, but PPB independent, divisions (Otegui & Staehelin, 2004; Brown & Lemmon, 2007). Furthermore, it is important to note that in most non-vascular plants, the PPB is absent or incomplete, or it is present in prophase only in certain cell types (Pickett-Heaps *et al*., 1999). From an evolutionary viewpoint, the PPB is thus a derived cytoskeletal structure that probably arose with the increased tissue complexity of vascular plants (Rasmussen *et al*., 2013).

What is the functional relevance of an apparent PPB-deficiency in dividing CSCs? One possibility is that the absence of a PPB is a way to conserve metabolic energy and to accelerate mitosis by avoiding the switch of cortical microtubules from transverse to longitudinal arrays in the very elongated CSCs. Considering its role in stabilizing spindle pole-to-pole axis perpendicular to the future division plane, the formation of a PPB in CSCs may also be unfavorable in light of possible spatial constraints. To test this hypothesis, it would be interesting to induce PPB formation in CSCs, for instance by ectopic expression of TRM7 and other potentially absent components of the microtubule organizing TTP complex (Spinner *et al*., 2013). In plant cells conducting PPB-independent cell divisions, such as gametophytic cells in non-vascular moss gametophore, a diffuse microtubule organizing center (MTOC) was detected during prophase. This acentrosomal microtubule cloud is dispersedly arranged and plays a crucial role in determining spindle and division plane orientation in cells dividing without prior PPB formation (Kosetsu *et al*., 2017). Also, in microspores and endosperm, radial microtubule arrays define division planes in the absence of PPBs (Krupnova *et al*., 2013). However, we did not detect an MTOC-like or any other specialized microtubule structure during CSC prophase, with the exception of the perinuclear microtubule array reminiscent of *trm678* mutants in root meristem cells. Since the antibody used for immunolabelling only detects α-Tubulin, further experiments to investigate the organization of γ-Tubulin could be conducted to confirm our observations. The molecular mechanisms underlying the variability in spindle orientation during metaphase and factors ensuring the robustness of CDZ component positioning remain unclear, but possibly involve the integration of mechanical cues in dividing CSCs. Further investigations of mechanisms underlying the control of division plane orientation including the mode of POK recruitment in CSCs will be essential to fully understand how tissue organization and radial growth are coordinated in the cambium.

## Acknowledgements

We thank David Brouchez (INRA Centre de Versailles-Grignon, France) for donating the *TRM7-3xYFP* line and the *trm678* triple mutant and Karin Schumacher (Centre for Organismal Studies, Heidelberg University, Germany) for donating the *pVHA-a3:VHA-a3-GFP* line. Work in TG’s group was supported by the Deutsche Forschungsgemeinschaft (DFG) through the FOR2581 consortium *Quantitative Morphodynamics of Plants* under grant number GR2104/6. Work in the group of SM was supported by DFG grant MU3133/6-2. XL was supported by a PhD completion grant of the *Landesgraduiertenförderung* program of the state Baden-Württemberg, a *heiDOCS* travel grant of the graduate academy of Heidelberg University, and a travel grant of the Schmeil foundation.

## Author contributions

XL and TG designed the study. XL performed experiments and analyzed data. LSS analyzed data. SM provided essential material and PL provided technical expertise. XL and TG wrote the manuscript with input from all authors.

## Data availability

The data that support the findings of this study are available in the supplementary material of this article. The code used for single cell transcriptome analysis is found at the GitHub repository [https://github.com/thomasgreb/Liu-et-al_CSC-divisions].

## Supporting Movies legends

**Movie S1 Maximum intensity projection of a discontinuous phragmoplast microtubule array in a CSC conducting a periclinal division.** Phragmoplast microtubule array was visualized by immunolabeling using an antibody against tubulin (green). Nuclei were stained by DAPI (blue). Slice space = 0.94 µm.

**Movie S2 Maximum intensity projection of a ring phragmoplast microtubule array in a CSC conducting an anticlinal division.** Phragmoplast microtubule array was detected by immunolabeling using an antibody against tubulin (green). Nuclei were stained by DAPI (blue). Slice space = 0.67 µm.

**Movie S3 Maximum intensity projection of a spindle microtubule array in a CSC conducting a periclinal division.** Spindle microtubule array was detected by immunolabeling using an antibody against tubulin (green). Nuclei were stained by DAPI (blue). Slice space= 0.62 µm.

**Movie S4 Maximum intensity projection of a spindle microtubule array in a CSC conducting an anticlinal division.** Spindle microtubule array was detected by immunolabeling using an antibody against tubulin (green). Nuclei were stained by DAPI (blue). Slice space= 0.60 µm.

**Movie S5 Maximum intensity projection of a perinuclear microtubule array in a prophase CSC.** The perinuclear microtubule array was detected by co-immunolabeling in a prophase CSC using an antibody against tubulin (green) and an antibody against the YFP protein (magenta). Nuclei were stained by DAPI (blue). The TRM7-3xYFP line was used. Slice space = 1 µm.

