## Supplemental Figure 1 for "Robust division orientation of cambium stem cells requires cortical division zone components but not the preprophase band"

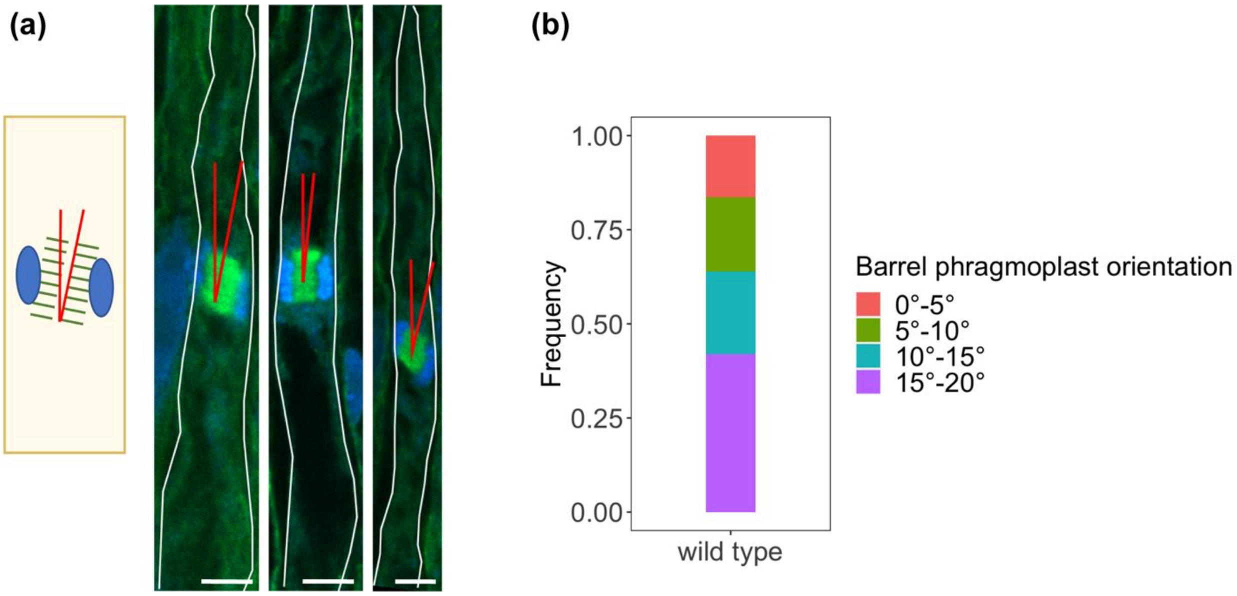


**Fig. S1 Disk phragmoplast is mobile in CSCs.** (a) Strategy to quantify disk phragmoplast orientation (red) in CSCs. Immunolabelling was performed using an antibody against tubulin (green). Nuclei were stained by DAPI (blue). Scale bar = 5 µm. (b) Quantification of disk phragmoplast orientation in wild type (n = 25).
