## Supplemental Figure 2 for "Robust division orientation of cambium stem cells requires cortical division zone components but not the preprophase band"

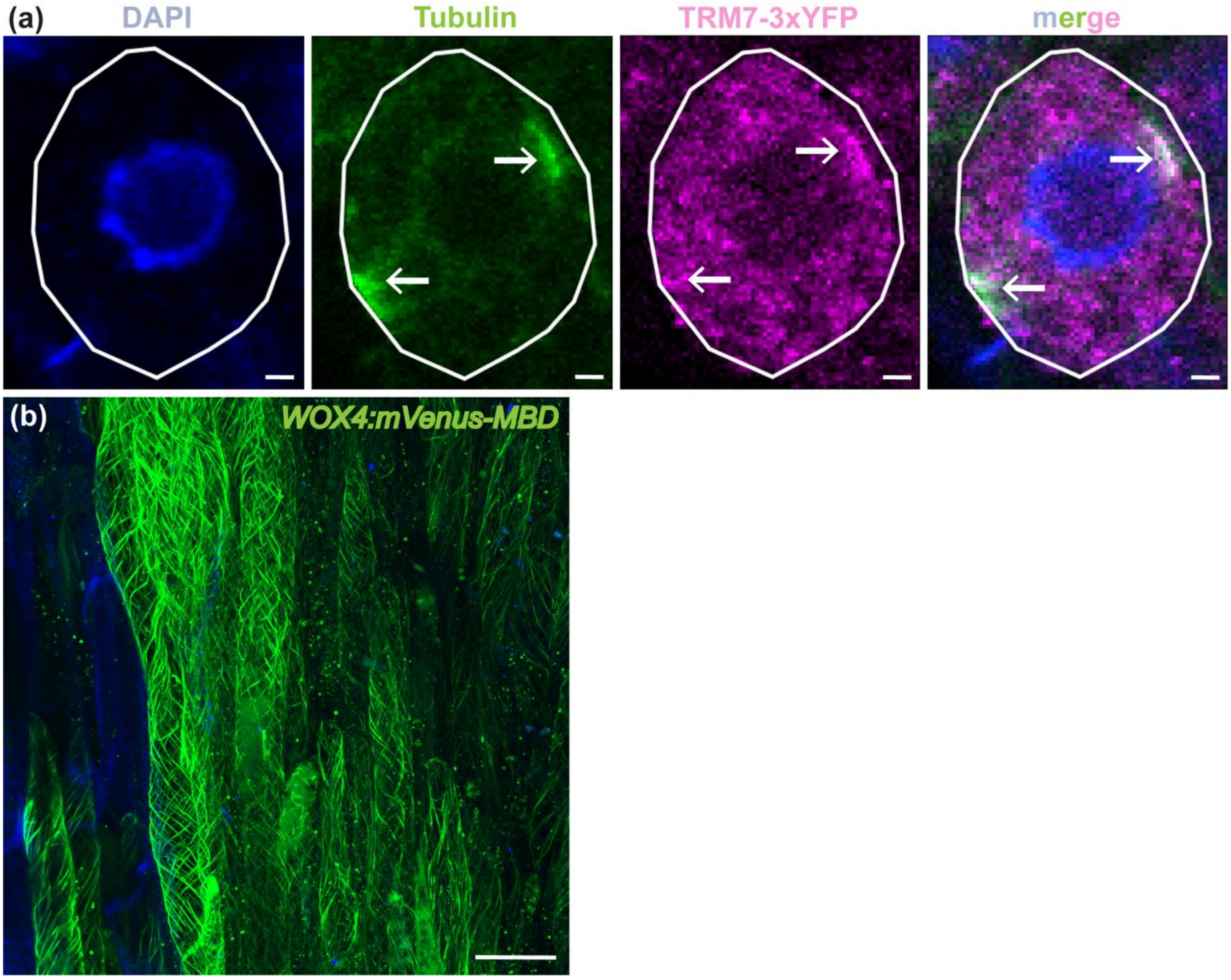


**Fig. S2 PPB microtubule arrays were detected in root tip cells and interphase cortical microtubules were detected in CSCs.** (a) PPB microtubule array was detected by co-immunolabeling in a dividing root tip cell using an antibody against tubulin (green) and an antibody against the YFP protein (magenta). Nuclei were stained by DAPI (blue). Cell boundaries are indicated by the white outline. Arrows point to the PPB microtubule arrays or TRM7-3xYFP-derived signals. The *TRM7-3xYFP* line was used in all analyses. n = 4 cells. Scale bar = 1 µm. (b) Max projections of signals from interphase cortical microtubules in CSCs visualized by live-cell imaging using the CSC-specific microtubule reporter *WOX4:mVenus-MBD* (green). Xylem autofluorescence was captured in blue. Scale bar = 20 µm.
