## Supplemental Figure 3 for "Robust division orientation of cambium stem cells requires cortical division zone components but not the preprophase band"

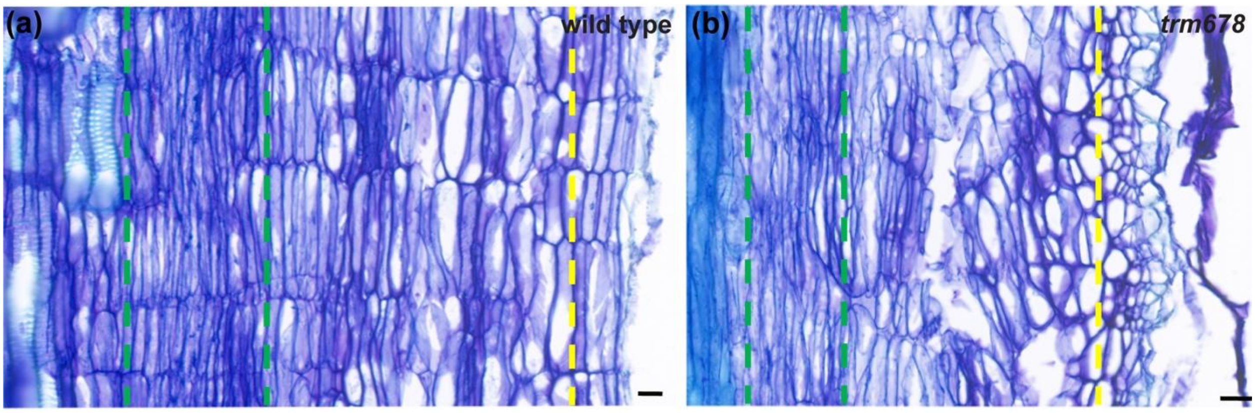


**Fig. S3 Toluidine bule-stained hypocotyl longitudinal sections from wild type and *trm678* mutants embedded in paraffin.** (a) wild type, (b) *trm678* mutant. Scale bar = 20 µm.
