## Supplemental Figure 4 for "Robust division orientation of cambium stem cells requires cortical division zone components but not the preprophase band"

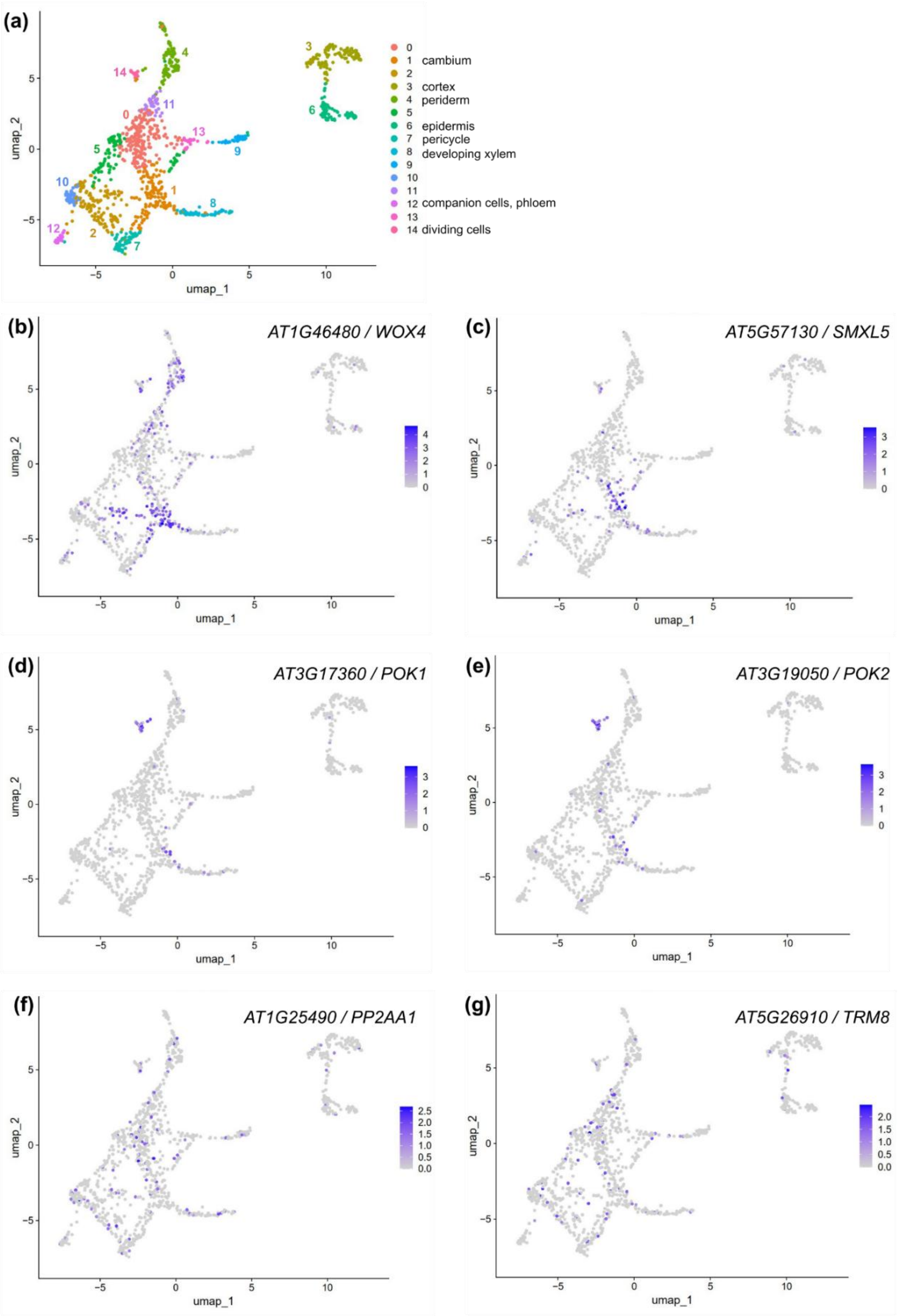


**Fig. S4 Analysis of CDZ-related gene expression in single cell transcriptome data described in Zhao et al., 2025.** (a) UMAP plot of snRNAseq data of 2,061 Arabidopsis hypocotyl nuclei organised in 14 clusters obtained through unsupervised clustering. (b-g) UMAP plot of transcript abundance of the two CSC marker genes *WOX4* and *SMXL5* (b, c), the CDZ-related genes *POK1* and *POK2* (d, e), and of the PPB-related genes *PP2A* and *TRM8*. Intensity of blue color indicate relative transcript abundance.
